# Design principles of Cdr2 node patterns in fission yeast cells

**DOI:** 10.1101/2023.04.19.537536

**Authors:** Hannah Opalko, Shuhan Geng, Aaron R. Hall, Dimitrios Vavylonis, James B. Moseley

## Abstract

Pattern forming networks have diverse roles in cell biology. Rod-shaped fission yeast cells use pattern formation to control the localization of mitotic signaling proteins and the cytokinetic ring. During interphase, the kinase Cdr2 forms membrane-bound multiprotein complexes termed nodes, which are positioned in the cell middle due in part to the node inhibitor Pom1 enriched at cell tips. Node positioning is important for timely cell cycle progression and positioning of the cytokinetic ring. Here, we combined experimental and modeling approaches to investigate pattern formation by the Pom1-Cdr2 system. We found that Cdr2 nodes accumulate near the nucleus, and Cdr2 undergoes nucleocytoplasmic shuttling when cortical anchoring is reduced. We generated particle-based simulations based on tip inhibition, nuclear positioning, and cortical anchoring. We tested model predictions by investigating Pom1-Cdr2 localization patterns after perturbing each positioning mechanism, including in both anucleate and multinucleated cells. Experiments show that tip inhibition and cortical anchoring alone are sufficient for the assembly and positioning of nodes in the absence of the nucleus, but that the nucleus and Pom1 facilitate the formation of unexpected node patterns in multinucleated cells. These findings have implications for spatial control of cytokinesis by nodes and for spatial patterning in other biological systems.

## Introduction

Spatial patterns are found in diverse biological systems, ranging from large ecosystems to intracellular structures. Many paradigms in pattern formation have emerged from embryonic development in multicellular organisms, but spatial patterns of signaling proteins also control pathways and processes in single cells (Karsenti, 2008; Green and Sharpe, 2015; Gross *et al*., 2017; McCusker, 2020; Zhu and Zernicka-Goetz, 2020; Mitchison and Field, 2021). Rod-shaped fission yeast cells (*S. pombe*) organize a set of cell cycle and cytokinesis proteins into a series of cortical “nodes,” which are nonmotile structures positioned in the cell middle (Paoletti and Chang, 2000; Morrell *et al*., 2004). The signals and organizing principles that position nodes in the cell middle to ensure proper cytokinesis are not fully understood.

Nodes are assembled and organized by the conserved protein kinase Cdr2, which binds the membrane through a KA1 domain (Martin and Berthelot-Grosjean, 2009; Moseley *et al*., 2009; Rincon *et al*., 2014). This domain also has clustering activity for node formation, and Cdr2 nodes then recruit downstream components such as anillin-like Mid1, cell cycle kinases Cdr1 and Wee1, and other proteins (Almonacid *et al*., 2009; Rincon *et al*., 2014; Guzmán-Vendrell *et al*., 2015; Allard *et al*., 2018). Two inhibitory signals from cell tips help to position Cdr2 nodes in the cell middle. First, the protein kinase Pom1 phosphorylates Cdr2 to inhibit its membrane-binding at tips (Martin and Berthelot-Grosjean, 2009; Moseley *et al*., 2009; Rincon *et al*., 2014). Second, membrane flows may limit Cdr2 node formation at growing cell tips (Gerganova *et al*., 2021). It is not clear if these inhibitory cues at cell tips are balanced by positive signals at the cell middle. Proper assembly of Cdr2 nodes also requires the GTPase Arf6, which localizes to nodes and serves as a cortical anchor (Opalko *et al*., 2022). Mutations that impair node formation and position lead to defects in cytokinesis and mitotic entry (Morrell *et al*., 2004; Martin and Berthelot-Grosjean, 2009; Moseley *et al*., 2009; Rincon *et al*., 2014; Guzmán-Vendrell *et al*., 2015), demonstrating the importance of understanding this spatial patterning system.

Here, we identified the nucleus as a positive signal for node positioning in the cell middle. We generated particle-based simulations of node formation and positioning by combined signals from the nucleus, Pom1, and Arf6. We tested elements of this model by moving or removing the cell nucleus, along with experiments using multinucleated cells that revealed unexpected striped patterns of nodes. Our combined experimental and modeling approach has revealed design principles that organize Cdr2 nodes for cell cycle progression and cytokinesis.

## Results and Discussion

### arf6 mutants reveal a connection between Cdr2 nodes and the nucleus

Prior work identified the GTPase Arf6 as a cortical anchor for Cdr2 nodes (Opalko *et al*., 2022). We observed Cdr2 in the nucleus of *arf6Δ* cells (Figure S1A). Nuclear localization of Cdr2 was enhanced by treatment of *arf6Δ* cells with leptomycin B (LMB) (Figures 1A-B and S1B), which inhibits Crm1-dependent nuclear export (Nishi *et al*., 1994; Kudo *et al*., 1998). LMB also induced minor accumulation of Cdr2 in the nucleus of wild type cells (Figures 1A-B). Thus, Cdr2 undergoes nuclear shuttling that is enhanced upon loss of cortical anchoring. Cdr2 associates with anillin-like node protein Mid1, which is known to undergo nuclear shuttling, and localization of Mid1 to interphase nodes requires Cdr2 (Paoletti and Chang, 2000; Moseley *et al*., 2009; Almonacid *et al*., 2009; Guzmán-Vendrell *et al*., 2015). We tested Cdr2 localization in the *mid1-nsm* mutant, which deletes Mid1 nuclear localization sequences and prevents Mid1 nuclear localization (Almonacid *et al*., 2009). Cdr2 did not localize in the nucleus of *mid1-nsm* cells, even when combined with *arf6Δ* and LMB treatments (Figures 1A-B). Once in the nucleus, Cdr2 only partially colocalized with Mid1 (Figure S1C). These results identify a previously unknown association between Cdr2 and the nucleus. The mechanism involves Cdr2 piggybacking into the nucleus with Mid1, counteracted by Cdr2 cortical anchoring mediated by Arf6.

**Fig 1:**
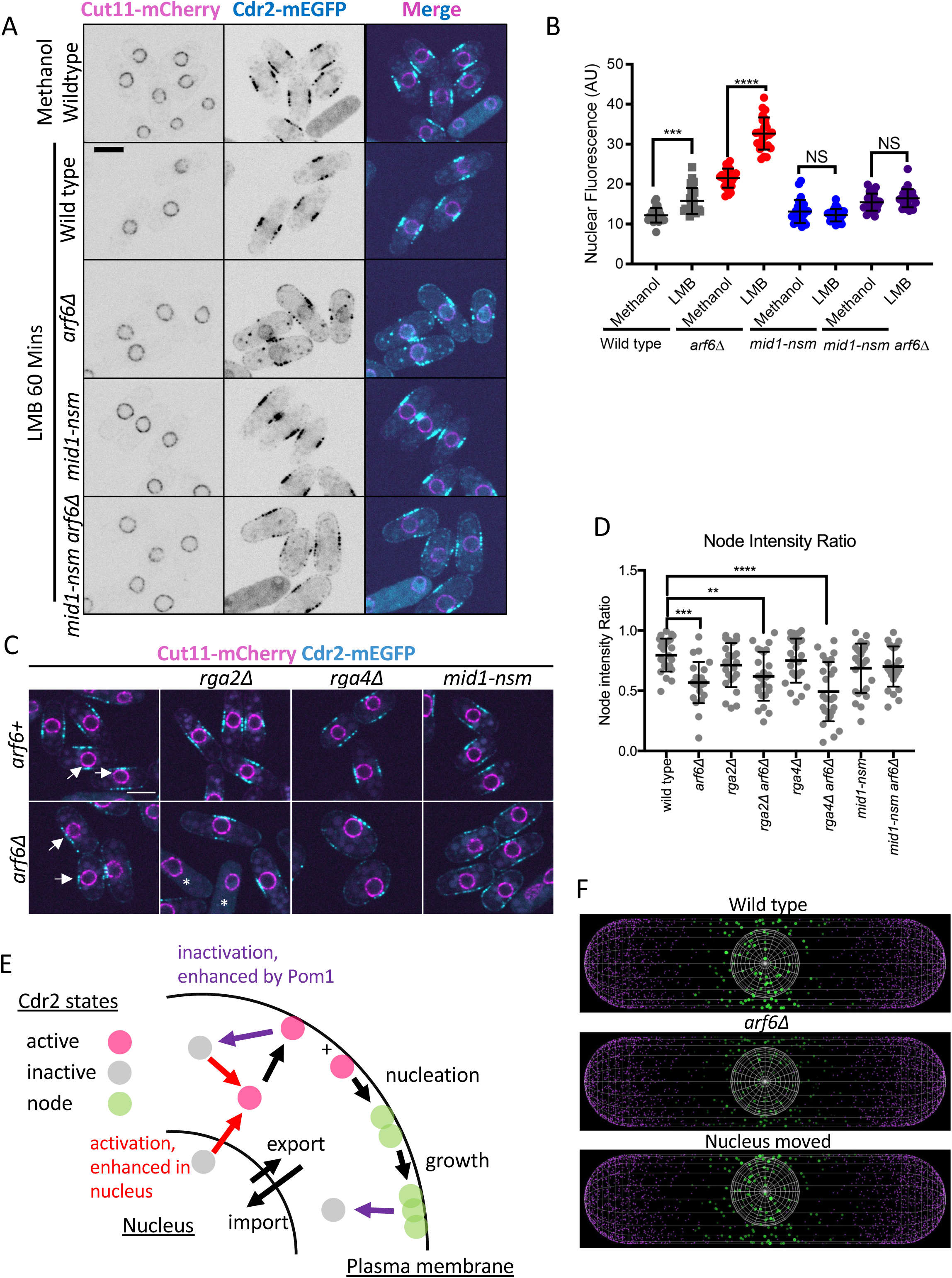
Regulated nuclear shuttling of Cdr2. (A) Single middle focal plane images of Cdr2 and the nuclear marker Cut11 in the indicated strains after leptomycin B (LMB) treatment. Scale bar, 5 μm. (B) Quantification of Cdr2 nuclear signal after LMB. n = 25 cells. ns >0.05, *** ≤0.001, **** ≤0.0001 by one-way ANOVA with Tukey’s multiple comparison test. (C) Single middle focal plane images of Cdr2 node localization in skinny (*rga2*Δ) and wide (*rga4*Δ) mutants. White arrows indicate off-center nuclei. Asterisks mark mitotic cells with diffuse cytoplasmic Cdr2. Scale bar, 5 μm. (D). (D) Quantification of Cdr2 node intensity ratio for the indicated strains. n > 25 cells. * ≤0.05, *** ≤0.001, **** ≤0.0001 by one-way ANOVA with Dunnett’s multiple comparison test. (E) Schematic of particle simulation model. Color of arrows indicates activation (red), inactivation (purple), or translocation (black). (F) Snapshots Cdr2 distribution after 50 min equilibration, for wild type, *arf6*Δ, and anucleate cell cases. Area and intensity of Cdr2 nodes (green) proportional to number of molecules per node. Pom1: magenta. Cell length: 11 μm.

We also observed a pronounced asymmetry in Cdr2 node positioning in cells lacking Arf6. More specifically, nodes were concentrated on one lateral side of *arf6Δ* cells (Figure 1C). The nucleus was pushed against the same lateral side, consistent with the nucleus acting as a local positive signal for node localization. In contrast, nodes were symmetrically distributed between the two sides of wild type cells even when the nucleus was pushed against one lateral side (Figure 1C), and nuclear positioning was not different between wild type and *arf6Δ* cells (Figure S1D-E). We measured the node intensity ratio between the two lateral cell sides (Figure S1E), which revealed a significant loss of node symmetry in *arf6Δ* cells (Figure 1D). We reasoned that decreasing cell width might suppress node asymmetry by reducing the distance between nucIei and cell sides, while increasing cell width might exacerbate asymmetry. Consistent with this prediction, node asymmetry in *arf6Δ* was partially rescued by the skinny mutant *rga2Δ* (Villar-Tajadura *et al*., 2008) and was enhanced by the wide mutant *rga4Δ* (Das *et al*., 2007) (Figures 1C,D). We conclude that the nucleus plays an instructive role in node positioning when cortical anchoring is reduced. This function appears to involve nuclear shuttling of Cdr2 because *arf6Δ* did not alter node symmetry in *mid1-nsm* mutants (Figures 1C,D), which prevent Cdr2 nuclear localization. These results indicate that the nucleus has local effects on node accumulation at membranes in concert with cortical tethering and cell tip inhibitors.

To understand the interplay between the nucleus, cortical tethering, and tip inhibitors in controlling node formation and localization, we generated particle-based simulations for Cdr2 molecules forming nodes in spatially restricted compartments of the plasma membrane (Figure 1E). In these simulations, Cdr2 molecules shuttle into and out of the nucleus. After leaving the nucleus, Cdr2 molecules are active to bind the plasma membrane, nucleate node assembly, and oligomerize during node growth. Activation in the nucleus was required to recapitulate the medial concentration pattern of nodes in cells (Figure S1F).

Node nucleation and growth are inhibited by Pom1 molecules localized in a cortical gradient concentrated at cell tips. Results from these simulations mimic experimental measurements of node number, size, and position (Figure S2). Altering model parameters allows us to simulate experimental perturbations. For example, we simulate the effects of *arf6Δ* on cortical tethering by increasing the off-rate of Cdr2 molecules (Figure 1F), based on prior photobleaching results (Opalko *et al*., 2022). Similarly, we simulated the effects of positioning the nucleus adjacent to the cell side in *arf6Δ* cells, which increased the local concentration of nodes similar to our experiments (Figure 1F, S1G). Based on the ability of this model to recapitulate our past experimental results, we used model-based predictions to test the role of the nucleus in node formation and patterning.

### Cdr2 nodes follow the nucleus

If the nucleus promotes Cdr2 node positioning, then moving the nucleus should impact nodes. We modeled this effect in simulations where the nucleus was displaced towards cell ends (Figure 2A). Modeling results showed accumulation of nodes near the displaced nucleus in a timeframe of 10-20 minutes (Figures 2B, S3A). These simulations showed a redistribution of Cdr2 from nodes near the cell middle towards new and existing nodes near the displaced nucleus, even though the node inhibitor Pom1 remained fixed at cell ends.

**Fig 2:**
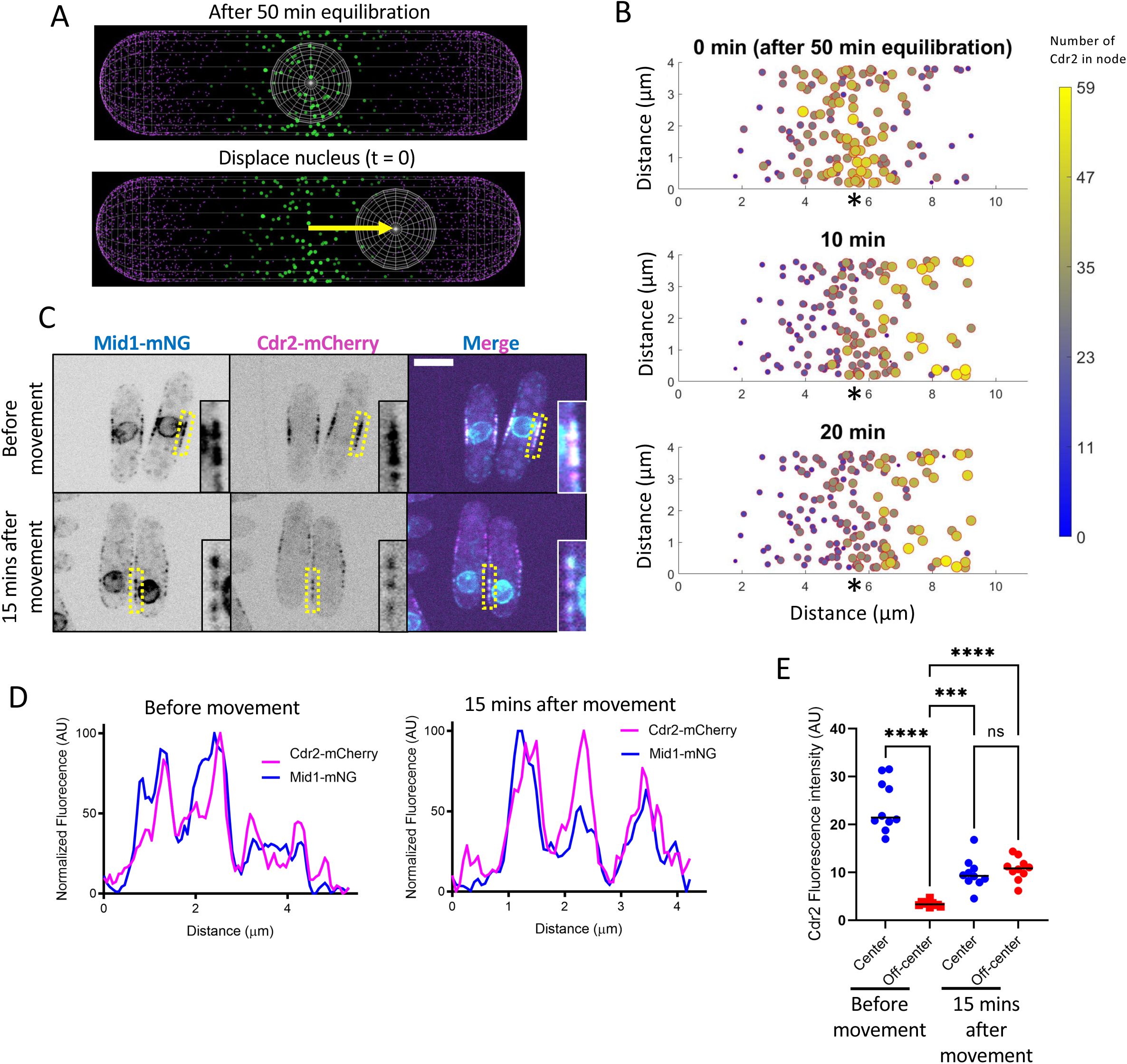
Cdr2 nodes follow displaced nuclei. (A) Start of nuclear displacement simulation after equilibrating Cdr2 for 50 min.. (B) Simulated response to nuclear movement with node area and color according to number of Cdr2 molecules. Asterisks mark the cell middle. (C) Single middle focal plane images of Cdr2 and Mid1 before and 15 minutes after nuclear displacement. Boxes show enlarged images of node region highlighted with a yellow dashed box. Strains are *cdc25-degron-DAmP* (*cdc25-dD*) mutation to increase cell length. Scale bar, 5 μm. (D) Line scans of the boxed regions in panel C of Cdr2 and Mid1 at the highlighted node region. Cdr2 and Mid1 colocalize in nodes before and after movement of the nucleus. (E) Quantification of Cdr2 fluorescence at the cell center or near the tip in cells before and after nuclear movement. N= 10 cells *** ≤0.001, **** ≤0.0001 by one-way ANOVA with Dunnett’s multiple comparison test.

We tested these predictions experimentally. The fission yeast interphase nucleus is positioned in the cell middle due to microtubules (Tran *et al*., 2001), which extend from the nuclear-embedded spindle pole body out to the cell ends. To displace nuclei, we treated cells with the microtubule depolymerizing drug MBC followed by centrifugation, leading to nuclear displacement caused by centrifugal force (Carazo-Salas and Nurse, 2006; Daga *et al*., 2006). Because the Cdr2-associating protein Mid1 has previously been shown to track with displaced nuclei (Daga and Chang, 2005), we monitored colocalization of Cdr2-mCherry and Mid1-mNG upon nuclear displacement. Prior to nuclear displacement, Cdr2 and Mid1 colocalized in nodes positioned exclusively in the cell middle (Figures 2C-E). After nuclear displacement, Cdr2 and Mid1 colocalized in nodes positioned over the displaced nucleus (Figures 2C-E). Importantly, we also observed nodes that remained in the cell middle (Figure 2E), as predicted by our simulations. We confirmed that Pom1 localizes to the cell ends in these experiments (Figures S3B-C), which means that Cdr2 node movement is not driven by strongly mislocalized Pom1. The presence of Pom1 at cell tips likely explains the absence of Cdr2 nodes at the extreme tips of cells even when the nucleus is nearby. However, we note the presence of a small but significant increase in Pom1 localization along cell sides in these experiments, which could promote Cdr2 dynamics and relocalization away from the cell center (Fig S3D). Together, our experimental and modeling results indicate that the cell nucleus provides a positive spatial cue for dynamic localization of nodes together with negative cues from Pom1 at cell ends.

### Node formation in anucleate cells

Given this novel role for the nucleus in positioning Cdr2 nodes, we next asked if cells can assemble and position nodes in the absence of a nucleus. We used centrifugation to displace mitotic nuclei, leading to the generation of anucleate and binucleate cells following division (Carazo-Salas and Nurse, 2006; Daga *et al*., 2006). Remarkably, we observed a medial band of Cdr2 nodes in newborn cells lacking a nucleus (Figure 3A, white asterisk). These nodes persisted for over 2 hours, during which time we also observed Pom1 at cell tips. To model this situation, we performed simulations for cells that lack a nucleus (Figure 3B). As in our experiments, these simulations showed the presence of medially positioned Cdr2 nodes based on the regulatory effects of Pom1 and cortical anchoring of Cdr2. The model also predicted a lower amount of Cdr2 molecules per node in the absence of nuclear Cdr2 regulation, as well as a reduced overall number of nodes (Figures S3E-F). We tested and confirmed both of these predictions experimentally for anucleate cells (Figure 3C-D). These results show that the nucleus promotes node formation but is not necessary for node assembly and positioning.

**Fig 3:**
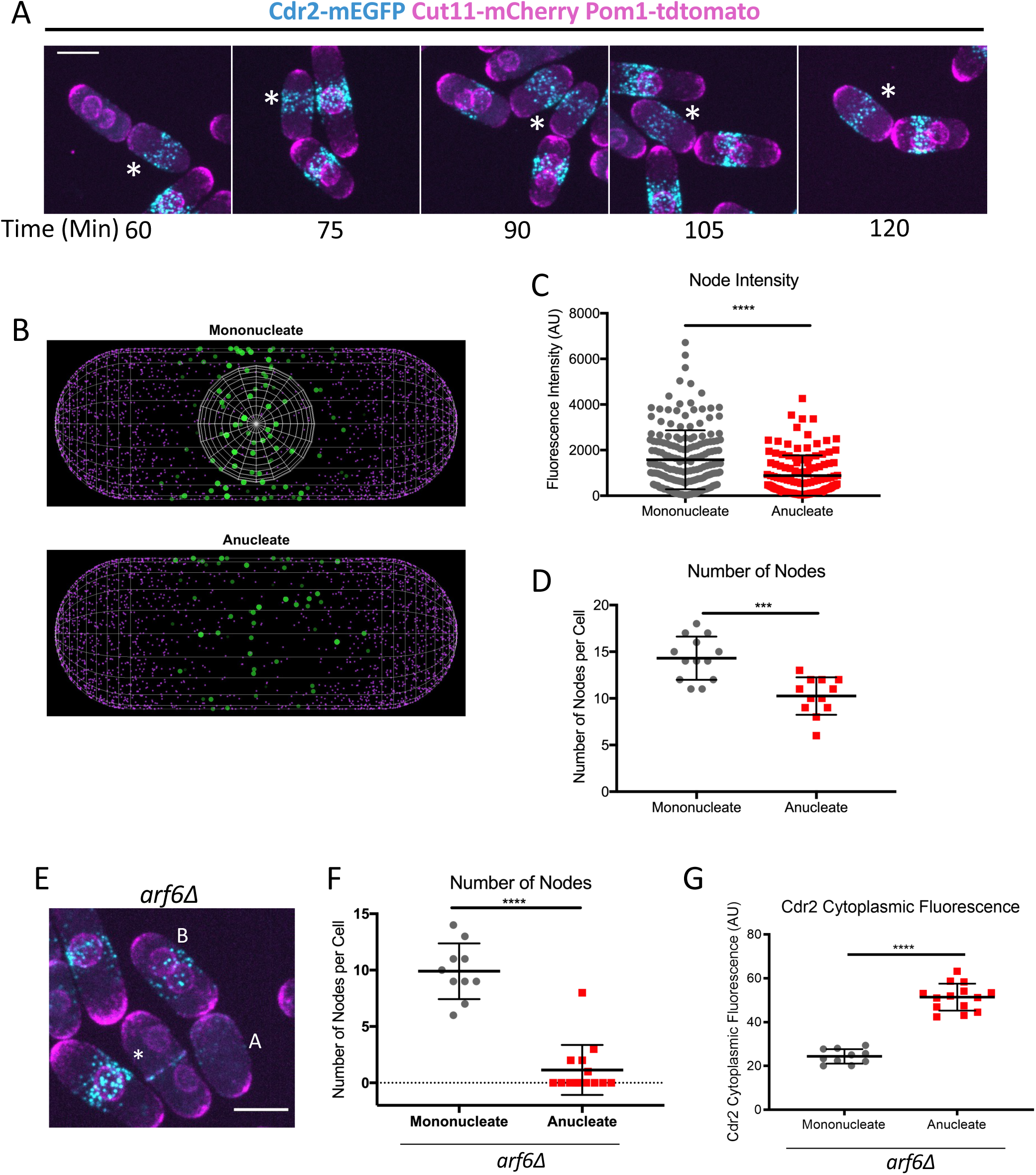
Formation of Arf6-dependent nodes in anucleate cells. (A) Maximum projection images of cells at the indicated timepoints after centrifugation. Images show anucleate and binucleate cells, with anucleate cell noted by the white asterisk. Cells are *cdc25-dD* to increase cell length. Scale bar 5 μm. (B) Simulated distribution of Cdr2 nodes for a mononucleate and anucleate cell after 50 min equilibration. (C) Fluorescence intensity of individual nodes 60 minutes after spin. n > 10 cells each. **** ≤ 0.0001. (D) The number of Cdr2 nodes per cell 60 minutes after spin. n > 10 cells. * ≤0.05, ** ≤0.01. (E) Arf6 is required for Cdr2 nodes in anucleate cells. Maximum projection image of cells 75 minutes after centrifugation. Anucleate cell labeled A, bincleate cell labeled B, dividing cell with diffuse cytoplasmic Cdr2 labeled with asterisk. Scale bar 5 μm. (F) The number of nodes per *arf6Δ* cell 60 minutes after spin. n > 10 cells. **** ≤0.0001. (G) Quantification of cytoplasmic Cdr2 levels in *arf6Δ* cells 60 minutes after spin. n > 10 cells. **** ≤0.0001. All statistical tests in this figure are Welch’s unpaired t-test.

Two additional experiments revealed regulatory mechanisms of nodes in anucleate cells. First, we tested the role of the cortical anchor Arf6, a positive regulator of nodes. Since Arf6 and the nucleus are both positive regulators of Cdr2 nodes in the cell middle, we reasoned that the dependence of nodes on Arf6 might be enhanced in anucleate cells. Consistent with this prediction, anucleate *arf6Δ* cells did not have Cdr2 nodes (Figures 3E-F), but instead displayed a higher cytoplasmic level of Cdr2 (Figure 3G). This result differs from the presence of nodes, albeit reduced in number and intensity, in mononucleated *arf6Δ* cells (Opalko *et al*., 2022). Simulations performed for anucleate *arf6Δ* cells also demonstrated a reduced number of Cdr2 nodes and intensity of Cdr2 per node (Figure S2F). We conclude that Cdr2 nodes can exist without either Arf6 or the nucleus, but removal of both positive regulatory mechanisms abolishes nodes in cells.

Second, we tested if Pom1 is required for node localization in the middle of anucleate cells. We confirmed that Pom1 forms a spatial gradient concentrated at the tips in anucleate cells (Figures S3G-H). Cdr2 nodes were asymmetrically positioned towards one cell tip in anucleate *pom1Δ* cells (Figure S3I-J), similar to effects in mononucleated *pom1Δ* cells (Martin and Berthelot-Grosjean, 2009; Moseley *et al*., 2009). This result means that Pom1 acts as a cell tip inhibitor of Cdr2 nodes both in the presence and absence of a nucleus. Interestingly, we found that the asymmetry of Cdr2 node localization in *pom1Δ* cells was partially suppressed in binucleated cells (Figures S3J), which indicates that doubling the positive regulatory signal provided by the nucleus in the cell middle can overcome loss of negative regulatory signals at cell tips. Together, these experiments reveal the nucleus is not required for Cdr2 nodes, but it reinforces node positioning in concert with other regulatory mechanisms including Arf6 and Pom1.

### Unexpected node patterns in multinucleated cells

In addition to testing how the absence of a nucleus impacts node patterning, we considered the opposite situation: multinucleated cells. Based on our results identifying the nucleus as a positive spatial cue for Cdr2 nodes, we predicted that nodes would either form separate bands over each nucleus in multinucleated cells, or alternatively nodes would form a large broad band covering all nuclei. To observe multinucleated cells, we generated an analog-sensitive allele of Sid1 kinase, which is required for fission yeast cytokinesis (Balasubramanian *et al*., 1998; Guertin *et al*., 2000). In the presence of analog drug 3-MB-PP1 to inhibit Sid1, these cells underwent synchronized nuclear divisions to contain 2, 4, and then 8 nuclei without intervening cytokinesis or septation (Figures S3K-L). In early binucleated stages, Cdr2-mEGFP formed a single, broad band of nodes over the two nuclei (Figure 4A). However, as these nuclei separated from each other, we observed a separate band of nodes over each nucleus. Following a brief stage of three Cdr2 node bands in early tetranucleated cells, we observed a surprising pattern of two bands of Cdr2 nodes in tetranucleated cells. These bands were positioned directly over the outer two nuclei, but Cdr2 nodes were absent from the inner two nuclei (Figure 4A). This unexpected pattern is consistent with a positive spatial cue from the outer two nuclei, but also reveals an additional signal that restricts nodes from the inner two nuclei.

**Fig 4:**
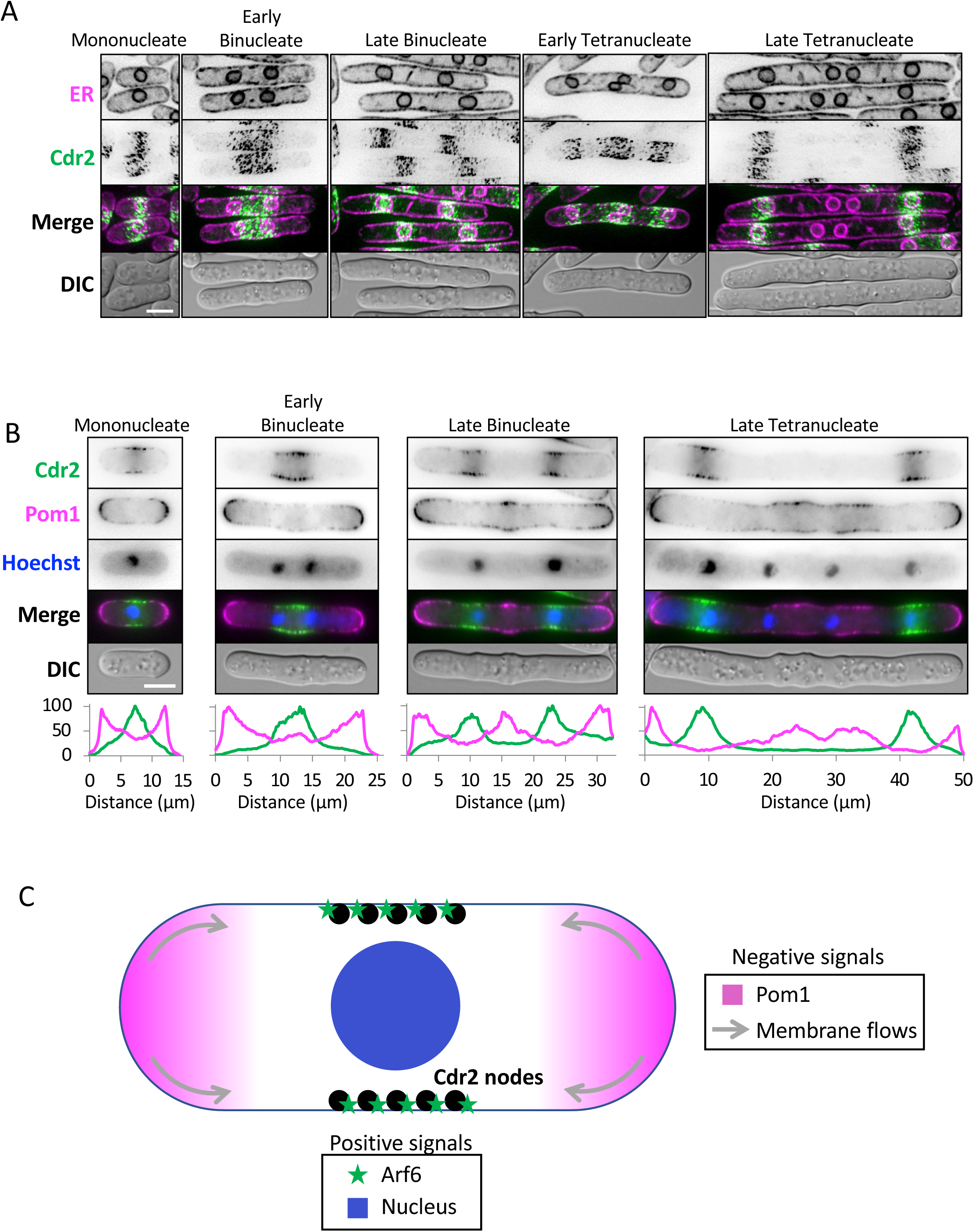
Spatial patterning of Cdr2 nodes in multinucleated cells. (A) Localization of Cdr2-mEGFP nodes in representative multinucleated cells. Nuclei and cell boundaries are marked by ER protein Sur4-mCherry. Images are maximum intensity projections of a Z-series. (B) Single middle focal plane images of multinucleated cells. Hoechst stain marks nuclei. Line scans of Cdr2 (green) and Pom1 (magenta) are shown below each image. Scale bars, 5 µm. (C) Working model for positioning of Cdr2 nodes by overlapping positive and negative signals.

We tested the possibility that Pom1 contributes to the unexpected spatial pattern of nodes in tetranucleated cells. We imaged Cdr2-mEGFP, Pom1-mCherry, and nuclei (Hoechst stain) in the same multinucleated cells (Figure 4B). Though restricted to cell ends in mononucleated cells and early binucleated cells, Pom1 was deposited in the middle of large binucleated cells, consistent with depletion of Cdr2 nodes from this site. Even more strikingly, Pom1 occupied the medial cortex of tetranucleated cells including the space directly over the inner two nuclei. In all cases, Pom1 was also present at the cell tips, where Cdr2 nodes were absent. We conclude that positive signals from nuclei combine with mutually exclusive localization of Pom1 and Cdr2, leading to banded node patterns in multinucleated cells.

## Conclusions

We combined experiments and simulations to generate a new model for how fission yeast cells mark their middle (Figure 4C). Our experiments revealed a role for the cell nucleus as a positive spatial cue for nodes. The organizing principles in patterning the Pom1-Cdr2 system resemble patterns observed during embryonic development (Briscoe and Small, 2015; Green and Sharpe, 2015; Schweisguth and Corson, 2019). Our work identifies the cell nucleus as a short-range activator for Cdr2 nodes, which are reinforced at the cell middle by Arf6-based cortical anchoring. These two positive signals are partially redundant as revealed by our anucleate cell experiments. Pom1 acts as an inhibitor to prevent node formation at cell tips (Martin and Berthelot-Grosjean, 2009; Moseley *et al*., 2009; Rincon *et al*., 2014). In addition, recent work showed that the flow of membrane lipids away from growing cell tips inhibits the stable tip localization of protein complexes such as nodes (Gerganova *et al*., 2021). Thus, robust spatial order in the fission yeast cell emerges from overlapping positive signals in the cell middle (nucleus and Arf6) combined with overlapping negative signals at cell tips (Pom1 and membrane flows).

Physical boundaries such as cell tips can influence pattern forming systems. Our experiments with multinucleated cells removed the boundary between each nucleus, revealing the potential for Pom1-Cdr2 to form complex patterns of node banding. These experiments also demonstrated that the inhibitor Pom1 overcomes local node activation over inner nuclei, consistent with previous work on ectopically targeted Pom1 (Martin and Berthelot-Grosjean, 2009; Moseley *et al*., 2009). Continued efforts combining experimental and modeling approaches are likely to uncover additional organizing similarities among seemingly divergent pattern-forming systems in biology.

## Acknowledgments

We thank Jace Curran for help in early stages of the modeling work, and members of the Moseley lab for discussions and comments on the manuscript. This work was supported by grants from the National Institutes of General Medical Sciences (NIGMS) to D.V. (R35GM136372) and to J.B.M. (R01GM099774 and R01GM133856).

## Materials and Methods

### Strain construction and media

Standard *S. pombe* media and methods were used (Moreno *et al*., 1991). Strains used in this study are listed in Table S1. Homologous recombination was used for C-terminal tagging and gene deletions (Bähler *et al*., 1998).

### Spinning disk confocal imaging

Live cell imaging in Figures 1-3 was performed using a Nikon Eclipse Ti-2E microscope with a Yokogawa CSU-WI spinning disk system and a Photometrics Prime BSI sCMOS camera. A 100x 1.45-NA CFI Plan Apochromat Lambda objective was used.

### Image analysis and quantification

ImageJ was used for all image analysis and quantification. All statistics and graphs were generated using Prism8 (Graphpad software).

For Leptomycin B treatment, cells were treated with 50ng/µl of leptomycin B or methanol for 60 minutes in EMM4S at 25°C. Cdr2 fluorescence was quantified by drawing a region of interest (ROI) around the nucleus based on Cut11-mCherry fluorescence. Cells were analyzed using a one-way ANOVA with Tukey’s multiple comparison test for statistical analysis.

For node intensity ratio, cells were grown in YE4S at 25°C. Cdr2 fluorescence was measured using the same-sized ROI surrounding the node region for each strain. At least 25 cells were analyzed using a one-way ANOVA with Dunnett’s multiple comparison test to compare each strain to the wildtype control. For nuclei placement, lines were drawn from the cell periphery using a brightfield image to the nuclear periphery using Cut11-mCherry as a marker. 25 cells were analyzed, and a one-way ANOVA with Tukey’s multiple comparison test was used for statistical analysis.

To make anucleate cells, cells were grown in EMM4S at 25°C. Cells were centrifuged at 15,000 rpm for 8 minutes in a microfuge, and then resuspended in EMM4S (Carazo-Salas and Nurse, 2006; Daga *et al*., 2006). Time indicated is time after centrifugation. The Pom1 gradient was measured through a sum projection of a timelapse of cells imaged every 5 seconds for 1 minute. A line was then used to trace one half of the cell, from one tip to the other tip. 10 cells were analyzed for mononucleate and anucleate cells. 10/10 mononucleate cells and 7/10 anucleate cells showed a clear Pom1 gradient. For node fluorescence intensity and node number per cell, z-series at 0.3 μm intervals were taken and the sum projections of the top half of the cell were used for quantification. ImageJ plugin comdetv5.5 was used to identify nodes. An ROI at the cell center was used and spot detection was set to a pixel size of 3 and an intensity threshold of 5. At least 10 cells were analyzed. To quantify Cdr2 cytoplasmic intensity, an ROI was drawn in the cytoplasm of middle single focal plane images. At least 10 cells were analyzed. For each analysis, a Welch’s unpaired t-test was used.

To move the nucleus, cells were grown in EMM4S at 25°C. Methyl-2-benzimidazole-carbamate (MBC) was added as previously described (Almonacid *et al*., 2009). In brief, 25 μg/mL of MBC was added to each culture for 5 minutes at 4°C. Cells were then centrifuged at 15,000 rpm for 8 minutes and then resuspended in EMM4S with MBC. For Figure 2D, Mid1 and Cdr2 fluorescence were each normalized by setting the maximum fluorescence value to 100 and the minimum value to 0 for each fluorophore. 11 cells were quantified for each condition. Pre-MBC treatment showed 11/11 cells to have colocalization between Cdr2 and Mid1. In MBC-treated condition 7/11 cells had colocalization between Cdr2 and Mid1, 4 cells had partial colocalization where we observed a Mid1 node that did not have Cdr2 fluorescence. For Figure 2E, Cdr2 fluorescence at the center and off-center was measured by drawing identically sized ROIs at the membrane positioned either in the cell center or off-center (near the tip) for 10 cells. This was done for cells imaged both before MBC addition and 15 minutes after MBC was added. Fluorescence was then background subtracted. A one-way ANOVA with Dunnett’s multiple comparison test was used to compare each column to all of the other column.

### Multinucleated cells

To generate analog-sensitive *sid1-as* cells, the *sid1+* gene including upstream promoter (800 nucleotides) and downstream terminator (500 nucleotides) were PCR amplified from genomic DNA and subcloned into integrative plasmid pJK148. Site-directed mutagenesis was used to introduce the M84G mutation resulting in plasmid pJM440 (sid1-as). A heterozygous diploid strain *sid1+/sid1Δ::kanMX6 leu1-32/leu1-32 ura4-D18/ura4-D18 ade6-M210/ade6-M216* was transformed with linearized pJM440. Transformants that grew on EMM4S-Leu were sporulated, the spores were separated by tetrad dissection, and *sid1Δ::kanMX6 leu1:[pJK148-sid1-as] ura4-D18 ade6-M21X* cells were selected by kanR and leu+ growth. To ensure even spacing of nuclei within the cytoplasm of multinucleated cells, *klp2-D25::ura4+* was introduced to these strains by genetic crosses.

The resulting *sid1-as* strains were grown in EMM4S media, and 10 µM 3-MB-PP1 was added to block sid1-as kinase activity. For imaging in Figure 4, cells were harvested by gentle centrifugation (1,000 rpm for 15 seconds in microfuge). Most of the supernatant was removed, and cells were resuspended in residual media containing inhibitor. Cells were imaged in liquid media under a coverslip with 0.4 µm step size with a personal DeltaVision Imaging System (Applied Precision/GE Healthcare), which included a customized Olympus IX-71 inverted wide-field microscope, a Photometrics CoolSNAP HQ2 camera, and an InsightSSI solid-state illumination unit. Images were acquired as Z-series with 0.4 µm step size and processed by iterative deconvolution in SoftWorx, followed by analysis in ImageJ. Images in Figure 4 show representative cells. The presented patterns were seen in all cells of a given category (n>20 each).

### Modeling Methods

Simulations were performed using Smoldyn version 2.67 (Andrews, 2017), a particle-based spatial stochastic simulator in which molecules are represented by point-like particles diffusing in a bulk or attached to surfaces. Their locations are updated and reactions between nearby particles occur at defined rates every timestep *dt*. In our model, the plasma membrane is represented as a cylinder of radius *R* and length *L* with two hemispherical caps of radius *R*. The nucleus is a spherical compartment of radius *R*_n_ centered in the cell. Based in part on previous modeling of Cdr2 nodes (Facchetti *et al*., 2019), we considered the simplest model for Cdr2 node formation via nucleation and growth, which incorporates activation near the nucleus and inhibition by Pom1. We did not model Pom1 gradient establishment, which has been addressed in earlier work (Saunders *et al*., 2012). Instead we placed *N*_Pom1_ stationary Pom1 particles on the plasma membrane, with uniform concentration along the cell tip hemispheres and with concentration declining exponentially with characteristic length 1 μm towards the middle of the cell, approximately matching the Pom1 number and distribution pattern in previous studies (Moseley *et al*., 2009; Rincon *et al*., 2014). We initialized *N*_Cdr2_ Cdr2 particles inside the nuclear compartment. Cdr2 diffusion and reaction (Figure 1F) were implemented as listed below. Input script for reference simulation is provided as a file in the Supplementary Materials.

#### Cdr2 states and activation

We assume Cdr2 particles can be in an active or inactive state in the cytoplasm or nucleus with diffusion coefficients *D*_cyt_ and *D*_nuc_, respectively. These two states represent in a simplified way how Cdr2 membrane binding and reactivity is influenced by phosphorylation (Moseley *et al*., 2009). Entry and exit of Cdr2 to the nucleus occurs with transmission coefficients *p*_nuc_out_ and *p*_nuc_in_, respectively. Activation of Cdr2 occurs when entering the nucleus or with a constant cytoplasmic background activation rate *k*^+^_bkg_. We further assume that only active Cdr2 can bind the plasma membrane and diffuse on it with diffusion coefficient *D*_mem._ In summary:

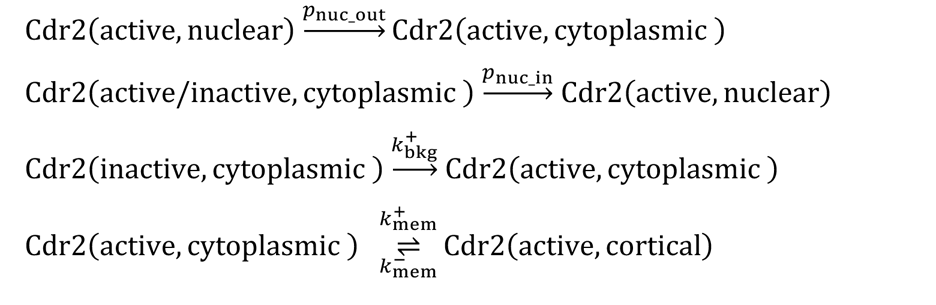

#### Cdr2 inactivation

Membrane bound active Cdr2 is deactivated and detached from the plasma membrane with a uniform constant rate *k*^−^_bkg_. Membrane bound active Cdr2 is additionally inactivated by Pom1 with rate constant *k*^−^_Pom1_:

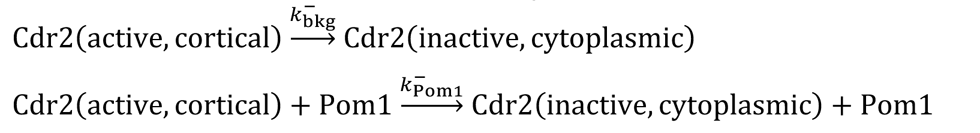

#### Node formation, growth, and disassembly

Two membrane-bound active Cdr2 molecules can react to form a node, represented as two stationary Cdr2 node particles placed on the same location on the plasma membrane, with rate constant *k*_nuc_. Additionally, active membrane-bound Cdr2 can accumulate in nodes by reacting with any Cdr2 in a node at rate constant *k*_grow_. A node consists of all Cdr2 node particles placed on the same location. Each Cdr2 in nodes dissociates into inactive cytoplasmic Cdr2 at rate *k*^−^_node_:

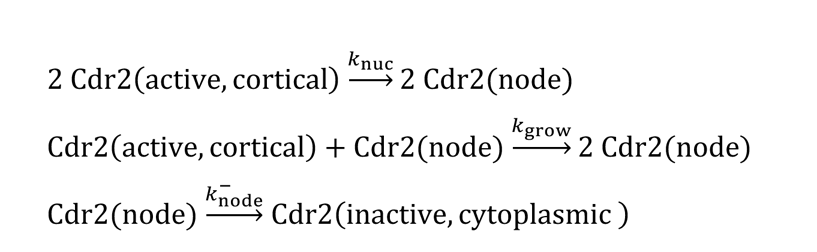

#### Equilibration and time step

We determined an appropriate equilibration time by measuring the node number and size distribution at different times. Node size grew rapidly once a few nodes were established and it slowly decreased as the total number of nodes gradually increased (Figure S2A). A time of 3000 s was sufficient to reach a stable state for the range of parameter values in the Table. The timestep *dt* was chosen to be large enough for computational efficiency after checking that smaller timesteps did not significantly improve accuracy.

#### Perturbations

In the anucleate condition, the nucleus was not included. For *arf6*Δ cells, we increased the node dissociation reaction rate constant since Arf6 acts as a cortical anchor to prolong nodal Cdr2 lifetime (Opalko *et al*., 2022). To investigate the effect of nuclear motion, the nuclear compartment and its contents were moved by 3 μm along the cell axis or 0.3 μm perpendicular to the cell axis after equilibration.

#### Parameter estimates

The plasma membrane attachment and detachment rate constants were assumed to be on the order of 1/s, faster than the Cdr2 node dissociation rate. The rate constant for membrane background deactivation, *k*^−^_bkg_, was also chosen to be of the same order. The rate constant of Pom1-mediated deactivation, *k*^−^_Pom1_, was large enough such that Cdr2 was excluded from cell tip under wild type conditions. These numbers provide a relative fraction of Cdr2 diffusing on the plasma membrane, nucleus and cytoplasm shown in Figure S2B. For the rate constants of node nucleation, *k*_*nuc*_, and growth, *k*_*grow*_, we performed a parameter scan to determine values that give realistic node number and size (Sayyad and Pollard, 2022) (Figure S2C-E). The background activation rate constant controls the relative strength of nuclear activation (the latter is simply determined by geometry in our model). We varied the background activation rate constant and selected a reference value that provides a significant reduction of Cdr2 in nodes in simulations of anucleate cells (Figure S2F).

#### Data analysis

The coordinates and state of each particle was periodically saved during the simulation and later processed in MATLAB to count the number of Cdr2 in the nucleus, in cytoplasm, on the cell membrane, and the number of nodes.

**Table 1.**
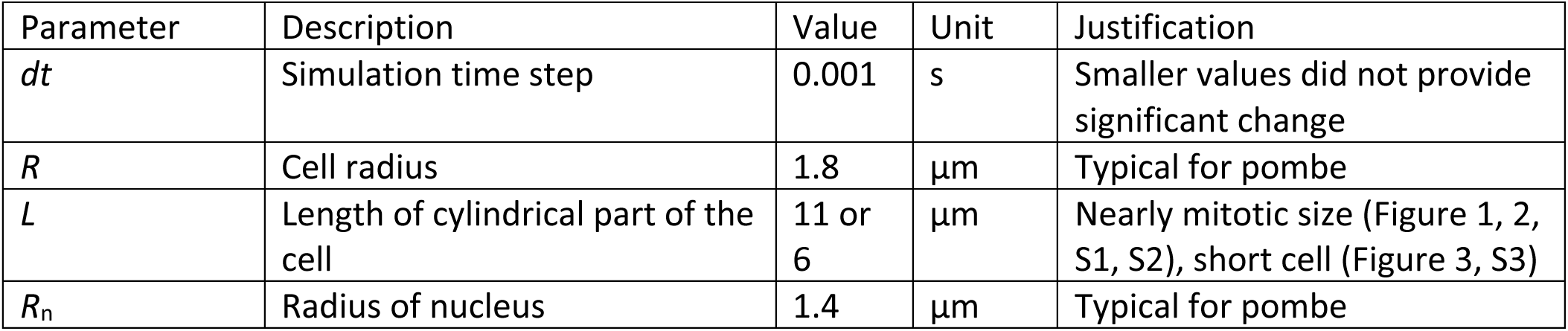
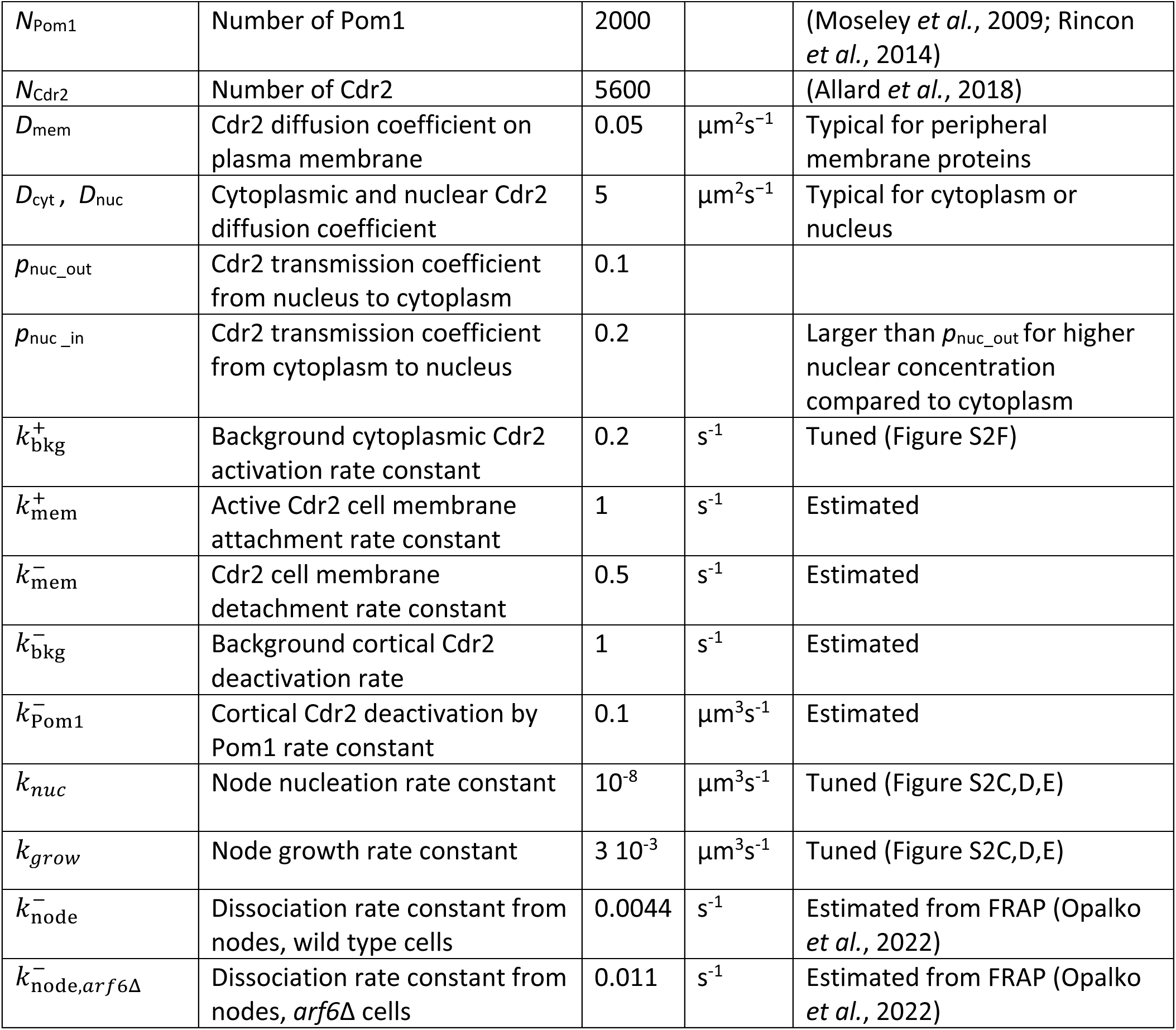
Simulation parameters.

---

**Supplemental Fig S1:** (A) Single middle focal plane images of Cdr2-mEGFP accumulation in the nucleus, marked by Cut11-mCherry, in *arf6Δ* cells. (B) Single middle focal plane images of Cdr2-mEGFP and Cut11-mCherry treated with methanol as control for LMB experiments. Scale bar, 5 μm. (C) Single middle focal plane images of Mid1-mNG and Cdr2-mCherry in the nucleus. Red boxed region is zoomed. (D) Quantification of nuclear position along the length of cells from the indicated strains. n > 25 cells. ** ≤0.01 by one-way ANOVA with Tukey’s multiple comparison test. (E) Schematic for nuclei placement and measuring node intensity ratio. (F) Snapshots of Cdr2 node distribution from simulations using different activation levels for the nucleus (N) or the cytoplasm (C). (G) Simulated response of Cdr2 in nodes to nuclear movement by 0.3 μm towards the top part of the cell, after a 50 min equilibration, for wild type and *arf6Δ* cases.

**Supplemental Fig S2:** (A) Node size distribution through a 10000s simulation, using wild type parameter values of Table 1. (B) Distribution Cdr2 at the end of 3000s simulation, using wild type parameter values of Table 1. (C-E) Dependence of equilibrated number of Cdr2 in nodes (panel C), number of nodes (panel D), and Cdr2 per node (panel E) on the values of node nucleation and growth rate constants. Other values same as in Table 1 for wild type cells. This parameter scan was used to determine reference values for node nucleation and growth. (F) Distribution of equilibrated number of Cdr2 per node in single cell simulations, with each dot representing one node. The background activation rate constant *k*^+^_bkg_ was changed for wild type, anucleate, *arf6*Δ, and *arf6*Δ anucleate cells. These simulations were used to tune the background activation constant such that removal of nucleus provides a significant reduction in number of Cdr2 in nodes.

**Supplemental Fig S3:** Analysis of cells with displaced or removed nuclei. (A) Histogram of simulated response of Cdr2 in nodes as function distance along the cell. (B) Pom1 remains concentrated at cell tips after nuclear displacement. Single middle focal plane images. Cells have *cdc25-dD* mutation to increase length. Scale bar 5 μm. (C) Linescan of fluorescence intensity for Cdr2-mEGFP and the combined Pom1-tdTomato and Cut11-mCherry along the length of cells before or after nuclear movement. (D) Pom1 levels along the lateral sides of cells is increased after nuclear displacement. *** ≤0.001 by Welch’s unpaired t-test. (E) Distribution of Cdr2 node size for two simulated cells after 50 min equilibration. (F) Number of Cdr2 nodes in mononucleate and anucleate cases for same cells as in E. (G) Pom1 gradient forms in anucleate cells. Sum projections of timelapse of Pom1. Yellow line depicts region for line scan in H. Scale bar, 5 μm.(F) Line scans of Pom1 gradient from each cell type. Yellow line from G depicts example line scan. (I) Multiple nuclei rescue Cdr2 localization defect in *pom1Δ* cells. Maximum project of a Z-series. Scale bar 5 μm. (J) Quantification of Cdr2 localization n>10 cells of each type done four times. (K) Percentage of cells with the indicated number of nuclei after addition of 3-MB-PP1 inhibitor to *sid1-as* cells. (L) Percentage of cells with the indicated number of nuclei after addition of DMSO control to *sid1-as* cells.

## References

1. Allard, CAH, Opalko, HE, Liu, K-W, Medoh, U, and Moseley, JB (2018). Cell size-dependent regulation of Wee1 localization by Cdr2 cortical nodes. J Cell Biol.

2. Almonacid, M, Moseley, JB, Janvore, J, Mayeux, A, Fraisier, V, Nurse, P, and Paoletti, A (2009). Spatial control of cytokinesis by Cdr2 kinase and Mid1/anillin nuclear export. Curr Biol 19, 961– 966.

3. Andrews, SS (2017). Smoldyn: particle-based simulation with rule-based modeling, improved molecular interaction and a library interface. Bioinformatics 33, 710–717.

4. Bähler, J, Wu, JQ, Longtine, MS, Shah, NG, McKenzie, A, Steever, AB, Wach, A, Philippsen, P, and Pringle, JR (1998). Heterologous modules for efficient and versatile PCR-based gene targeting in Schizosaccharomyces pombe. Yeast 14, 943–951.

5. Balasubramanian, MK, McCollum, D, Chang, L, Wong, KC, Naqvi, NI, He, X, Sazer, S, and Gould, KL (1998). Isolation and characterization of new fission yeast cytokinesis mutants. Genetics 149, 1265–1275.

6. Briscoe, J, and Small, S (2015). Morphogen rules: design principles of gradient-mediated embryo patterning. Development 142, 3996–4009.

7. Carazo-Salas, RE, and Nurse, P (2006). Self-organization of interphase microtubule arrays in fission yeast. Nat Cell Biol 8, 1102–1107.

8. Daga, RR, and Chang, F (2005). Dynamic positioning of the fission yeast cell division plane. Proceedings of the National Academy of Sciences 102, 8228–8232.

9. Daga, RR, Lee, K-G, Bratman, S, Salas-Pino, S, and Chang, F (2006). Self-organization of microtubule bundles in anucleate fission yeast cells. Nat Cell Biol 8, 1108–1113.

10. Das, M, Wiley, DJ, Medina, S, Vincent, HA, Larrea, M, Oriolo, A, and Verde, F (2007). Regulation of cell diameter, For3p localization, and cell symmetry by fission yeast Rho-GAP Rga4p. Mol Biol Cell 18, 2090–2101.

11. Facchetti, G, Knapp, B, Flor-Parra, I, Chang, F, and Howard, M (2019). Reprogramming Cdr2-Dependent Geometry-Based Cell Size Control in Fission Yeast. Curr Biol 29, 350–358.e4.

12. Gerganova, V, Lamas, I, Rutkowski, DM, Vještica, A, Castro, DG, Vincenzetti, V, Vavylonis, D, and Martin, SG (2021). Cell patterning by secretion-induced plasma membrane flows. Sci Adv 7, eabg6718.

13. Green, JBA, and Sharpe, J (2015). Positional information and reaction-diffusion: two big ideas in developmental biology combine. Development 142, 1203–1211.

14. Gross, P, Kumar, KV, and Grill, SW (2017). How Active Mechanics and Regulatory Biochemistry Combine to Form Patterns in Development. Annu Rev Biophys 46, 337–356.

15. Guertin, DA, Chang, L, Irshad, F, Gould, KL, and McCollum, D (2000). The role of the sid1p kinase and cdc14p in regulating the onset of cytokinesis in fission yeast. EMBO J 19, 1803–1815.

16. Guzmán-Vendrell, M, Rincon, SA, Dingli, F, Loew, D, and Paoletti, A (2015). Molecular control of the Wee1 regulatory pathway by the SAD kinase Cdr2. J Cell Sci 128, 2842–2853.

17. Karsenti, E (2008). Self-organization in cell biology: a brief history. Nat Rev Mol Cell Biol 9, 255– 262.

18. Kudo, N, Wolff, B, Sekimoto, T, Schreiner, EP, Yoneda, Y, Yanagida, M, Horinouchi, S, and Yoshida, M (1998). Leptomycin B inhibition of signal-mediated nuclear export by direct binding to CRM1. Exp Cell Res 242, 540–547.

19. Martin, SG, and Berthelot-Grosjean, M (2009). Polar gradients of the DYRK-family kinase Pom1 couple cell length with the cell cycle. Nature 459, 852–856.

20. McCusker, D (2020). Cellular self-organization: generating order from the abyss. Mol Biol Cell 31, 143–148.

21. Mitchison, TJ, and Field, CM (2021). Self-Organization of Cellular Units. Annu Rev Cell Dev Biol 37, 23–41.

22. Moreno, S, Klar, A, and Nurse, P (1991). Molecular genetic analysis of fission yeast Schizosaccharomyces pombe. Meth Enzymol 194, 795–823.

23. Morrell, JL, Nichols, CB, and Gould, KL (2004). The GIN4 family kinase, Cdr2p, acts independently of septins in fission yeast. J Cell Sci 117, 5293–5302.

24. Moseley, JB, Mayeux, A, Paoletti, A, and Nurse, P (2009). A spatial gradient coordinates cell size and mitotic entry in fission yeast. Nature 459, 857–860.

25. Nishi, K, Yoshida, M, Fujiwara, D, Nishikawa, M, Horinouchi, S, and Beppu, T (1994). Leptomycin B targets a regulatory cascade of crm1, a fission yeast nuclear protein, involved in control of higher order chromosome structure and gene expression. J Biol Chem 269, 6320–6324.

26. Opalko, HE, Miller, KE, Kim, H-S, Vargas-Garcia, CA, Singh, A, Keogh, M-C, and Moseley, JB (2022). Arf6 anchors Cdr2 nodes at the cell cortex to control cell size at division. J Cell Biol 221, e202109152.

27. Paoletti, A, and Chang, F (2000). Analysis of mid1p, a protein required for placement of the cell division site, reveals a link between the nucleus and the cell surface in fission yeast. Mol Biol Cell 11, 2757–2773.

28. Rincon, SA et al. (2014). Pom1 regulates the assembly of Cdr2-Mid1 cortical nodes for robust spatial control of cytokinesis. J Cell Biol 206, 61–77.

29. Saunders, TE, Pan, KZ, Angel, A, Guan, Y, Shah, JV, Howard, M, and Chang, F (2012). Noise reduction in the intracellular pom1p gradient by a dynamic clustering mechanism. Dev Cell 22, 558–572.

30. Sayyad, WA, and Pollard, TD (2022). The number of cytokinesis nodes in mitotic fission yeast scales with cell size. Elife 11, e76249.

31. Schweisguth, F, and Corson, F (2019). Self-Organization in Pattern Formation. Dev Cell 49, 659– 677.

32. Tran, PT, Marsh, L, Doye, V, Inoué, S, and Chang, F (2001). A mechanism for nuclear positioning in fission yeast based on microtubule pushing. J Cell Biol 153, 397–411.

33. Villar-Tajadura, MA, Coll, PM, Madrid, M, Cansado, J, Santos, B, and Pérez, P (2008). Rga2 is a Rho2 GAP that regulates morphogenesis and cell integrity in S. pombe. Mol Microbiol 70, 867– 881.

34. Zhu, M, and Zernicka-Goetz, M (2020). Principles of Self-Organization of the Mammalian Embryo. Cell 183, 1467–1478.

