## Supplemental Table S1 for "Design principles of Cdr2 node patterns in fission yeast cells"

Table S1. Yeast strains used in this study

| | Genotype | Source |
|---|---|---|
| JM1043 | sid1Δ::kanMX6 sid1-as::leu1+ cdr2-mEGFP::kanMX6 pom1-tdTomato::natR klp2-D5::ura4+ ura4-D18 ade6-M21X | This study |
| JM1132 | sid1Δ::kanMX6 sid1-as::leu1+ cdr2-GFP::natR sur4-mCherry::natR klp2-D5::ura4+ ura4-D18 ade6-M21X | This study |
| JM7477 | cdr2-mEGFP::kanMX6 cut11-mCherry::kanMX6 rga2Δ::kanMX6 | This study |
| JM7478 | cdr2-mEGFP::kanMX6 arf6Δ::natR cut11-mCherry::kanMX6 rga2Δ::kanMX6 | This study |
| JM7497 | cdr2-mEGFP::kanMX6 arf6Δ::natR cut11-mCherry::kanMX6 rga4Δ::kanMX6 | This study |
| JM7505 | cdr2-mEGFP::kanMX6 mid1::ura4+ leu+ :Pmid1-mid1nsm leu1-32 cut11-mcherry::kanR | This study |
| JM7509 | arf6Δ::hphR cdr2-mEGFP::kanMX6 mid1::ura4+ leu+ :Pmid1-mid1nsm leu1-32 cut11-mcherry::kanR | This study |
| JM7511 | rga4Δ::kanMX6 cdr2-mEGFP::KanMX6 cut11-mcherry::KanMX6 | This study |
| JM7592 | cdr2-mEGFP::kanMX6 cut11-mCherry::kanMX6 pom1-tdtomato::nat cdc25-degron-DAmP::kanMX6 ura4-D18 | This study |
| JM7606 | cdr2-mEGFP::kanMX6 cut11-mCherry::kanMX6 pom1Δ::natR cdc25-degron-DAmP::kanMX6 | This study |
| JM7695 | cdr2-mEGFP::KanMX6 cut11-mcherry::kanMX6 | This study |
| JM7696 | cdr2-mEGFP::KanMX6 cut11-mcherry::kanMX6 arf6Δ::natR | This study |
| JM7707 | cdr2-mEGFP::kanMX6 cut11-mCherry::kanMX6 arf6Δ::natR pom1-tdtomato::natR h? | This study |
| JM7769 | mid1-yomNeonGreen::HphR cdr2-mCherry::natR cdc25-degron-DAmP::kanMX6 | This study |
