## Supplementary figures and images for "Design principles of Cdr2 node patterns in fission yeast cells"

### Figure S1

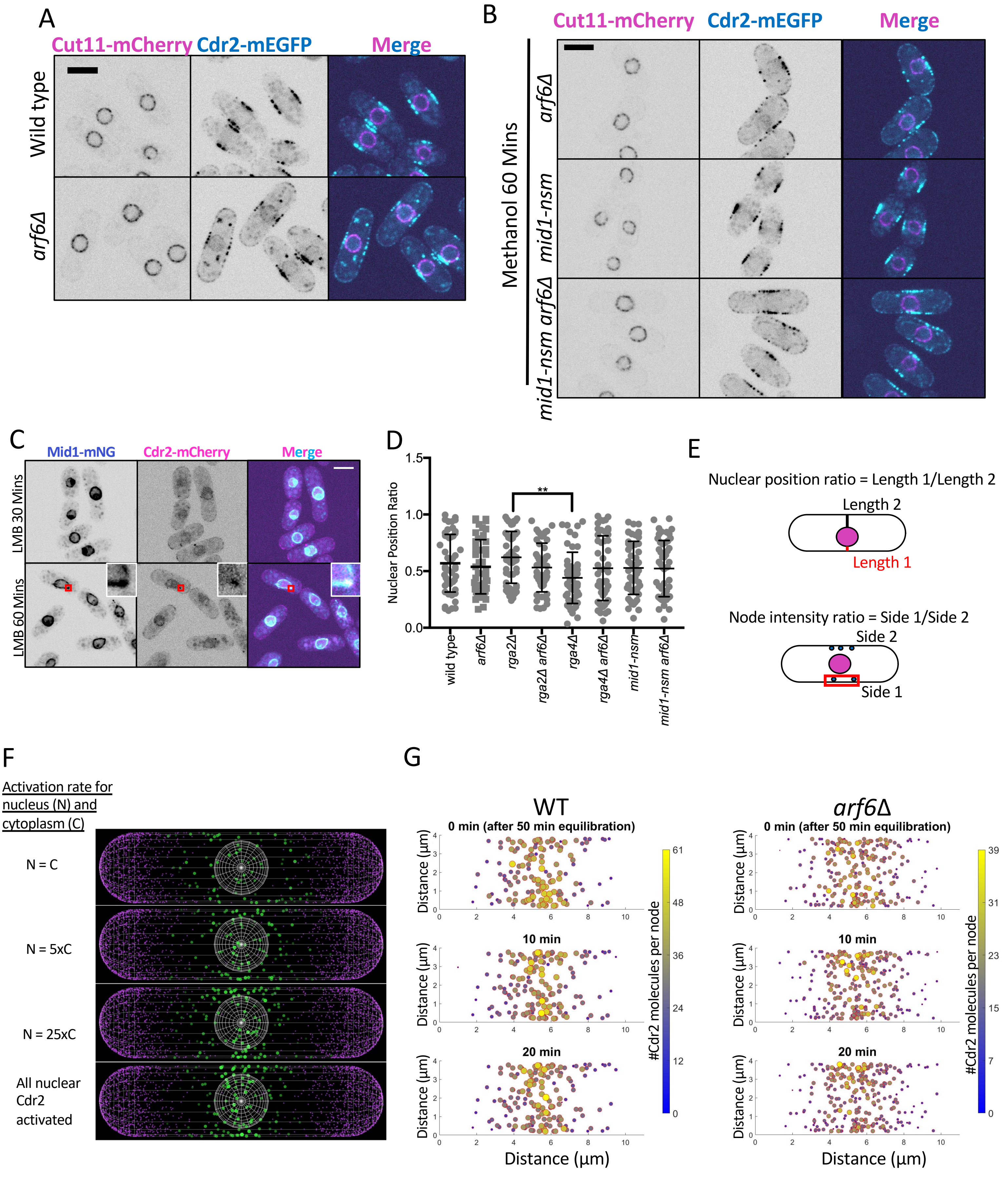


Supplemental Figure S1

### Figure S2

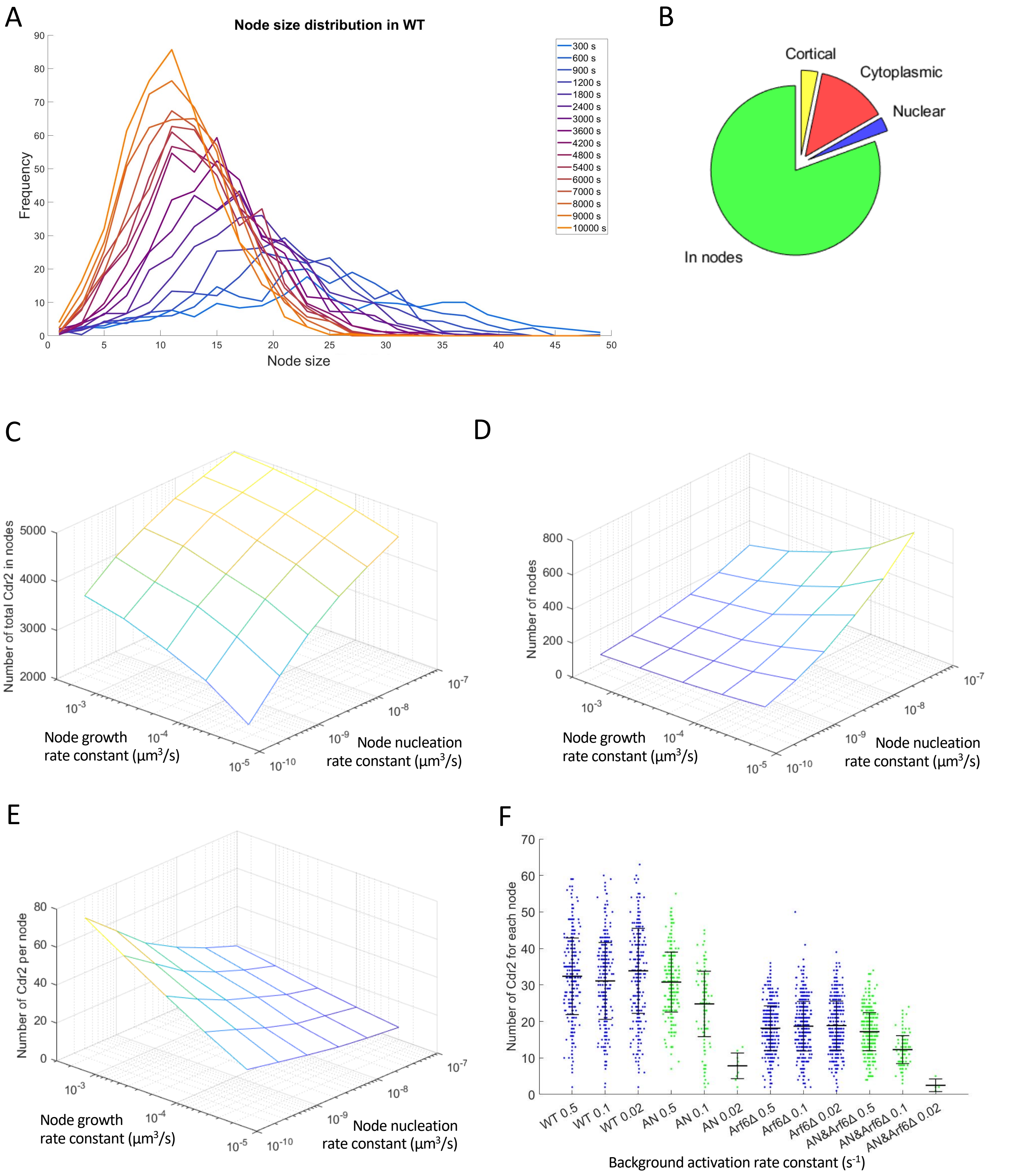


Supplemental Figure S2

### Figure S3

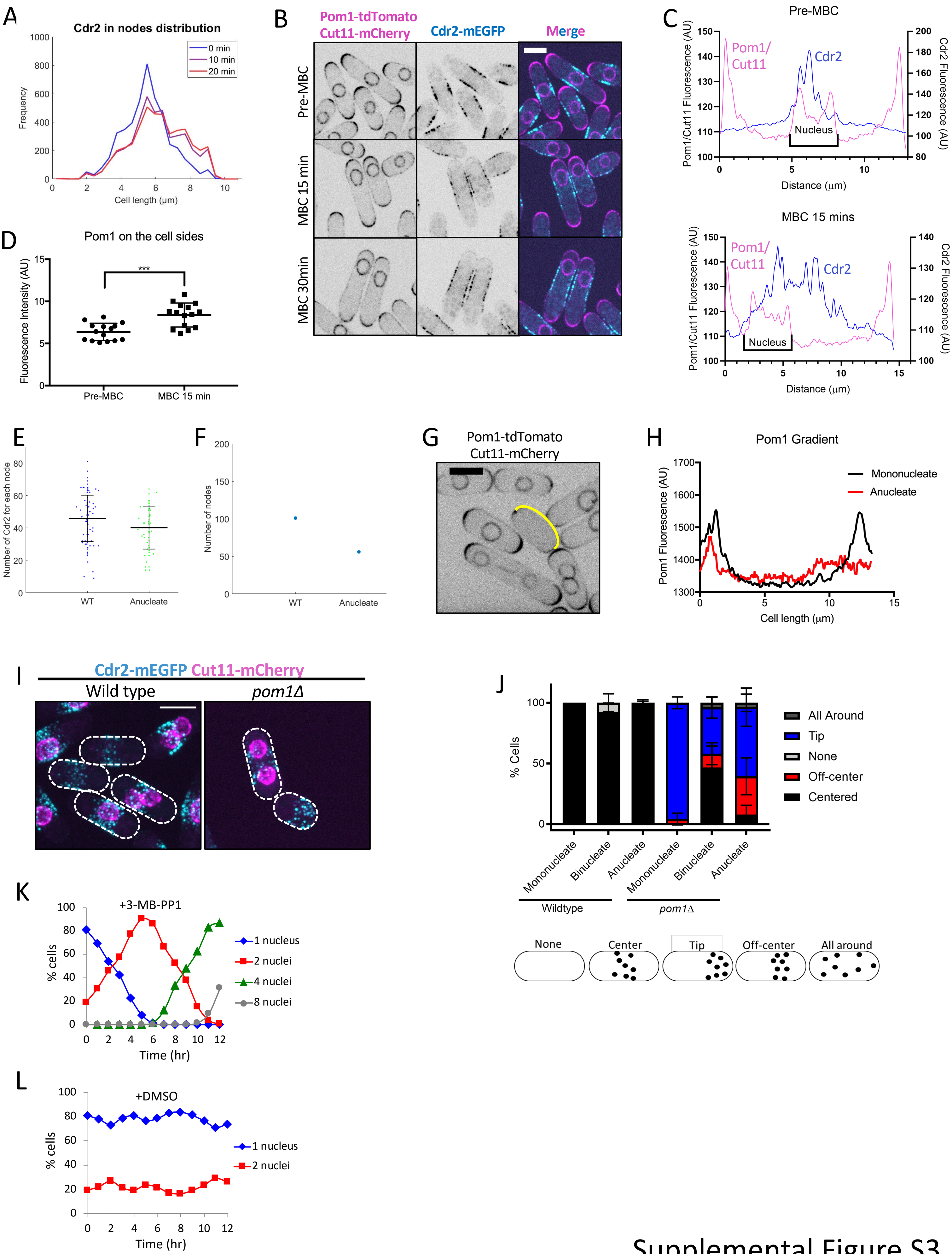


Supplemental Figure S3
